# Antiviral activity of Tecovirimat against Mpox virus clades 1a, 1b, 2a and 2b

**DOI:** 10.1101/2024.12.20.629622

**Authors:** Jeanne Postal, Florence Guivel-Benhassine, Françoise Porrot, Quentin Grassin, Jack M. Crook, Riccardo Vernuccio, Valérie Caro, Jessica Vanhomwegen, Pablo Guardado-Calvo, Etienne Simon-Lorière, Laurent Dacheux, Jean-Claude Manuguerra, Olivier Schwartz

## Abstract

The recent Mpox virus (MPXV) outbreak was caused by a novel and more pathogenic clade 1b virus. We compared the antiviral efficacy of Tecovirimat in cell culture, against the two ancestral clades 1a and 2a, the clade 2b that circulated in 2022, and the recent clade 1b virus. We report that Tecovirimat similarly inhibits the replication of all four MPXV clades, at nanomolar concentrations (nM). Our results suggest that Tecovirimat remains a therapeutic option against the latest clade 1b virus.

---

The zoonotic *Orthopoxvirus* Mpox virus (MPXV) includes two main clades (1 and 2) relevant to human transmission ^1^. Two major MPXV outbreaks occurred since 2022 ^1 2 3^ and were declared public health emergencies of international concern (PHEIC) by the WHO in July 2022 and August 2024, respectively. The first outbreak was caused by a clade 2b strain that quickly spread worldwide, resulting in about 100,000 cases and 200 deaths ^3^. The second outbreak was due to a novel and more pathogenic clade 1b strain. As of December 2024, this upsurge resulted in more than 55,000 reported or suspected cases and about 1,000 deaths in the Democratic Republic of Congo (DRC) and neighboring countries including Burundi, Rwanda and Uganda ^4^. A few imported clade 1b cases have also been reported in Europe, USA, Canada and Thailand ^5^. Prevention measures include patient isolation and care, and vaccines, which have started to be distributed in some African countries in fall 2024.

Tecovirimat (TPOXX^TM^) has been widely used as an antiviral treatment ^6 7^. Tecovirimat was initially approved by the US Food and Drug Administration as an anti-smallpox molecule. It is authorized in Europe and UK for the treatment of Mpox and other orthopoxviruses. In the US, 7,000 individuals with Mpox have received Tecovirimat ^6^. Tecovirimat binds to the conserved Orthopox protein F13L, that elicits production of wrapped virions from infected cells ^8^. Tecovirimat acts as a molecular glue, triggering F13L dimerization and blocking its function^8^. Tecovirimat is active against MPXV lethal challenge in monkey models. In humans, Tecovirimat is safe but there are divergent reports of clinical efficacy ^6 7^. A shorter time to symptom recovery in treated patients has been observed in some real-world studies but not in others ^7^. The PALM007 placebo-controlled trial was initiated in DRC in 2022, before the emergence of the novel clade 1b ^9^. Preliminary results did not show reduction in time to Mpox skin lesion resolution in Tecovirimat treated patients ^7 9^. Similarly, interim results of the STOMP trial, performed in South America, USA and Asia did not demonstrate Tecovirimat efficacy in skin and mucosal lesion resolution’s time, in patients with mild to moderate Mpox ^10^. Most of the patients received Tecovirimat more than 5 days after symptom onset ^10^.

About 24 Tecovirimat-resistant mutations have been reported in the F13L protein ^7 8^ (Appendix pages 4-6). These mutations are present in 1% of sequenced viruses in USA ^11^. Some of the mutations were observed in chronically MPXV-infected immunocompromised patients treated with Tecovirimat for long periods of time ^11^. Tecovirimat-resistant MPXV strains have also been occasionally detected in non-treated patients, indicating that the resistant viruses are transmissible ^7^.

The antiviral activity of Tecovirimat against MPXV clade 1b has not been characterized. We compared the sensitivity of the recent clades 1b and 2b to Tecovirimat in cell culture. The clade 1b strain was isolated in August 2024 in Sweden from a traveler returning from Africa ^5^. The clade 2b virus was isolated in France in 2022 ^12^. We included the ancestral clades 1a and 2a strain as controls. We used U2OS cells as targets, as they are highly sensitive to MPXV infection ^13^ (Appendix page 2). Tecovirimat efficiently inhibited the four viral strains, with similar EC50s, ranging from 15.3 to 25.8 nM (Figure and Appendix page 3). These EC50s correspond to those reported for the clade 2b strain ^14^. Moreover, an analysis of 310 available sequences of the Clade 1b virus did not identify known Tecovirimat-associated mutations in the F13L protein, whereas they were detected at low frequency among the larger number of Clade 2b genome sequences (10,968 sequences, Appendix pages 5-8).

Our *in vitro* results may not directly translate into clinical efficacy. However, combined with the known antiviral activity of the molecule in animal models, our results suggest that Tecovirimat remains a therapeutic option for the recently circulating clade 1b virus. Ongoing and future trials in different countries should help determining the optimal timing of administration and other parameters that may impact efficacy.

*O.S. and P.G.C. have a patent application for Poxvirus immunogens not used in the present study (PCT/EP2024/063801). The remaining authors declare no competing interests*.

**Figure.**
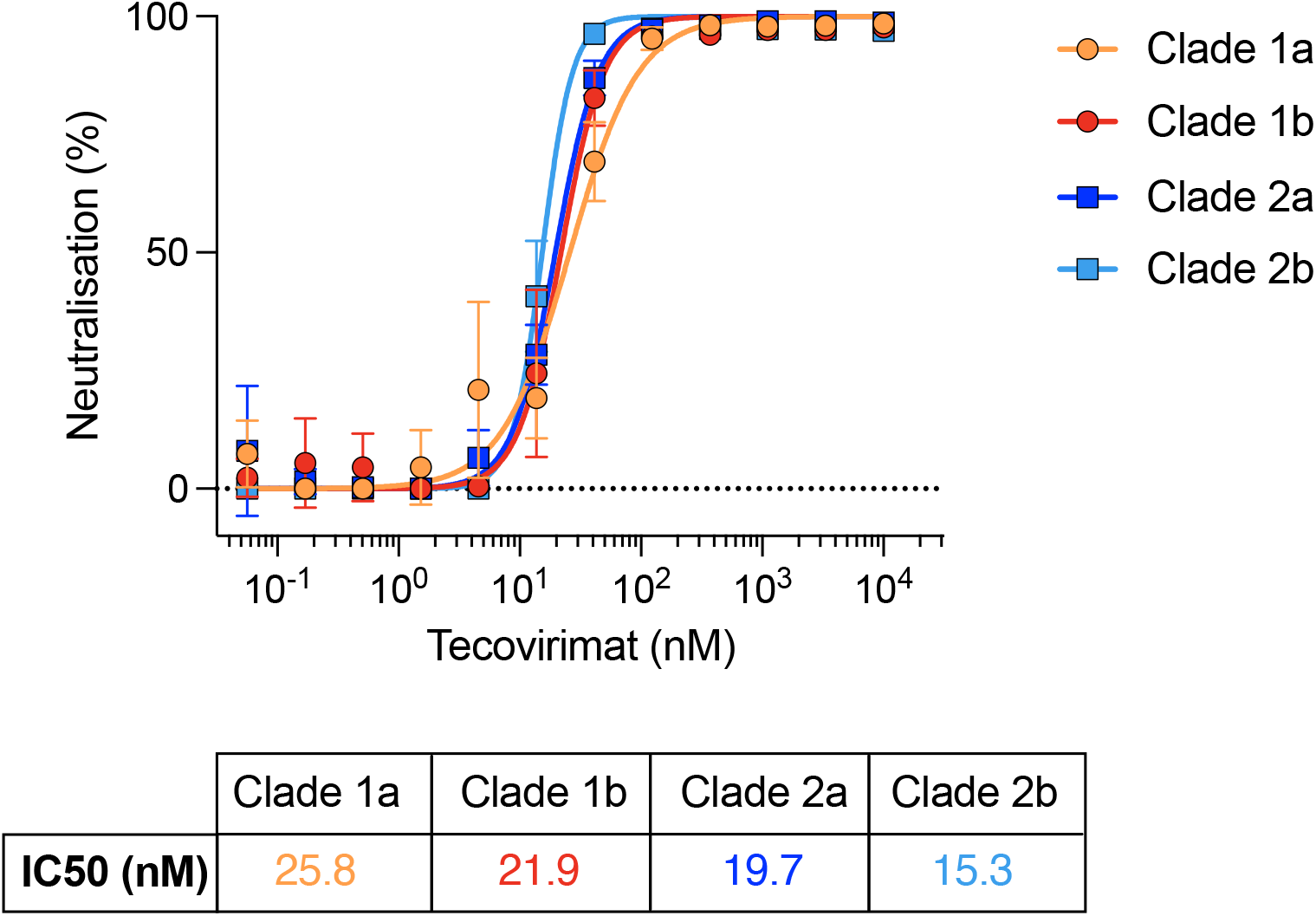
Comparison of the antiviral activity of Tecovirimat against different clades of MPXV. Dose-response analysis of inhibition of the indicated MPXV clades by Tecovirimat. Data are presented as mean ± standard deviation (s.d.) of triplicates from one experiment representative of at least 3 independent experiments. The IC50 values for Tecovirimat are indicated in the lower panel.

