## Supplemental data for "Antiviral activity of Tecovirimat against Mpox virus clades 1a, 1b, 2a and 2b"

Table of content

### Supplemental Figures and Tables

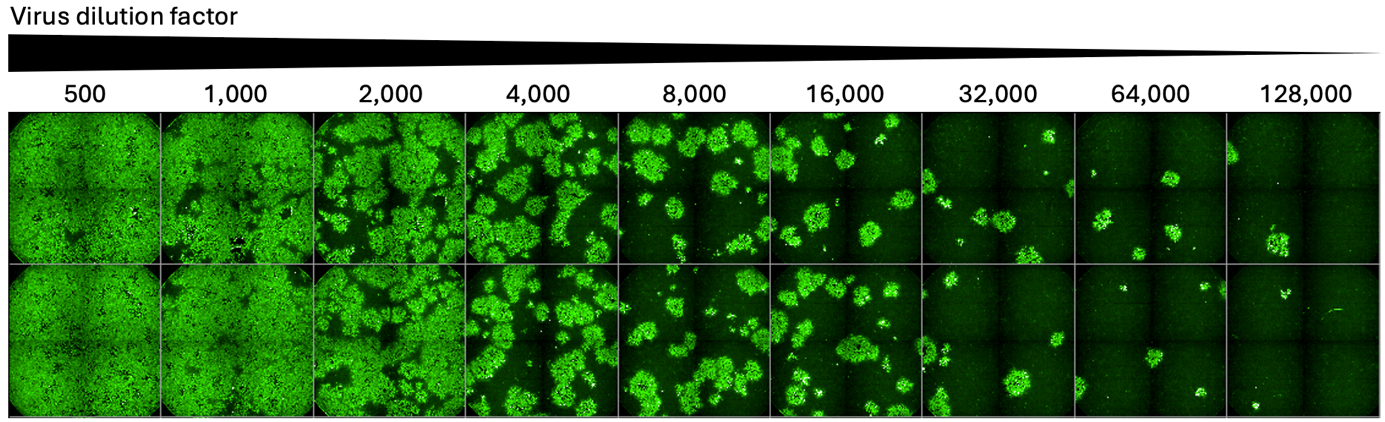

#### **Figure S1. Titration of MPXV clade 1b in U2OS cells.**

Indicated dilutions of the MPXV clade 1b viral stock ^1^, initially amplified in Vero cells, were incubated on U2OS cells in duplicates. After 48h, cells were fixed and MPXV was stained using a cross-reactive polyclonal anti-VACV antibody. Infected cells formed foci of different size, reflecting the known direct cell-cell transmission of poxvirus ^2^. The MPXV^+^ areas were quantified. The number of infected cells correlated with the viral inoculum. We selected a non-saturating inoculum for Tecovirimat inhibition experiments (1:8,000 in the depicted image). One representative experiment, with each well in duplicate, is shown. Each square represents an independent well. MPXV strains from other clades were similarly titrated.

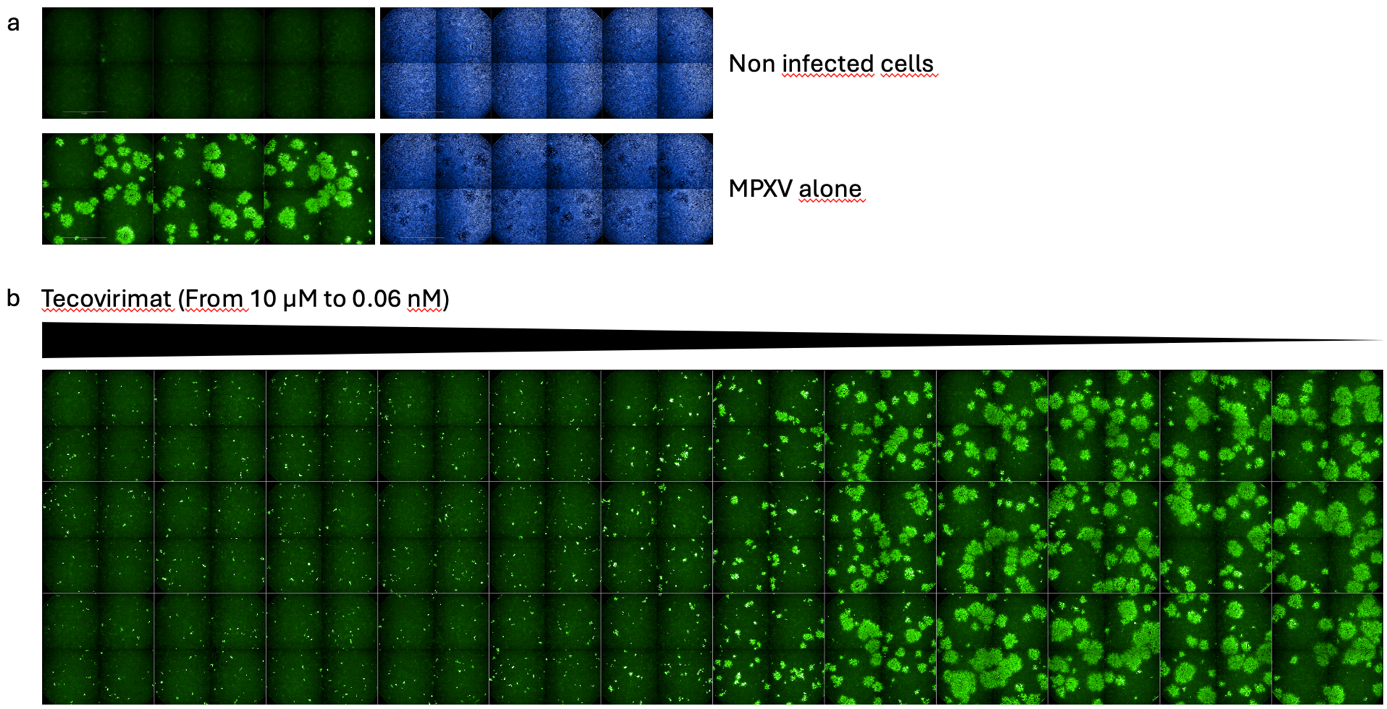

#### **Figure S2. Inhibition of MPXV clade 1b by Tecovirimat.**

a. Example of non-infected and MPXV-infected cells, as described in Figure S1. Left panels: After 48h, cells were fixed and MPXV was stained using a cross-reactive polyclonal anti-VACV antibody, to visualize infected cells (in green). Right panels: Cells were stained with Hoechst 33342 to visualize nuclei (in blue). Each square represents an independent well.

b. U2OS cells were preincubated for 4 h with the indicated doses of Tecovirimat and infected with MPXV clade 1b. After 48h, cells were fixed and MPXV was stained using a cross-reactive polyclonal anti-VACV antibody. The MPXV^+^ areas were quantified and the percentage of inhibition of infection was calculated. One representative experiment, with each well in triplicate, is shown. Each square represents an independent well. The small clusters of infected cells at high concentrations of Tecovirimat likely reflects the mode of action of the drug, which acts at a late step of the viral life cycle. The antiviral effect of Tecovirimat on other MPXV strains was similarly analyzed.

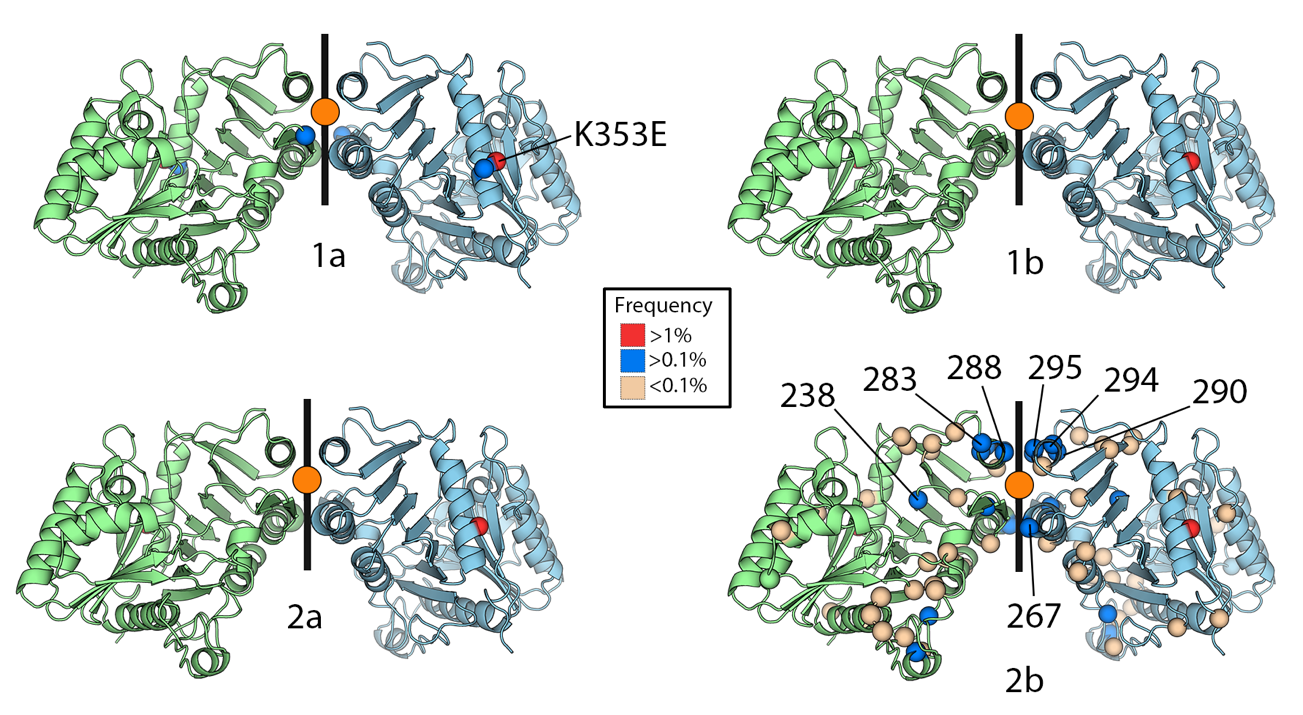

#### **Figure S3. Representation of the F13L protein from the four MPXV clades.**

The structure of a dimeric F13L protein was adapted from Vernuccio ^3^. One protomer is colored blue and the other green. The 2-fold symmetry axis is indicated by a black line, and the Tecovirimat binding site is marked by an orange-filled circle. Colored spheres highlight the positions and frequencies of observed mutations within different lineages, based on data available in the *GISAID EpiPox* database. The most common mutation, K353E, is located outside of the Tecovirimat binding site and is not associated with Tecovirimat resistance. Amino acid residues known to confer Tecovirimat resistance are labeled. Detailed information on the resistance impact of each mutation and their frequencies can be found in Tables S1 and S2, respectively.

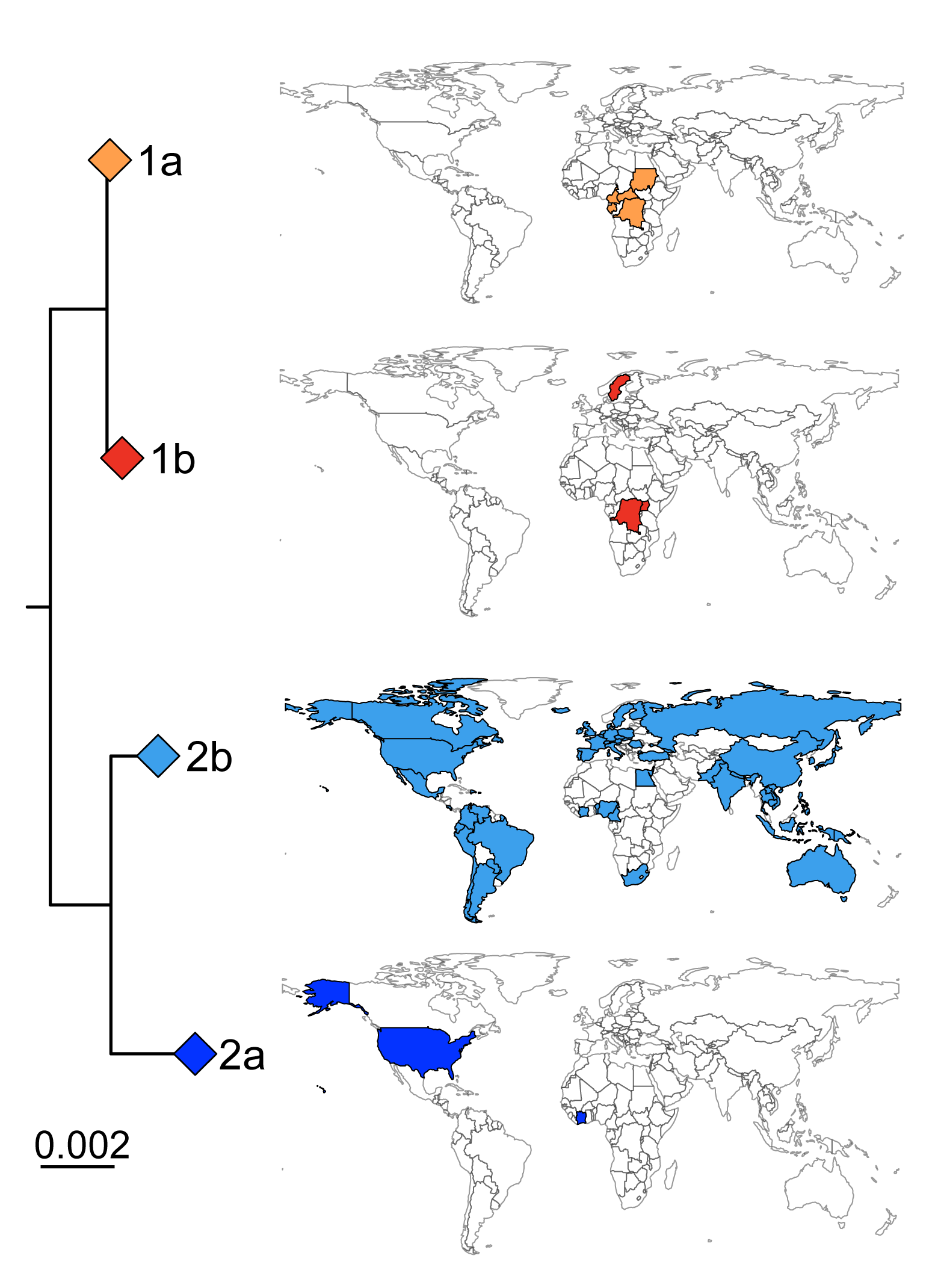

#### **Figure S4. Phylogenetic representation of the MPXV clades from which F13L genes were extracted for the mutational analysis depicted in Table S2.**

Genome sequences were retrieved from GISAID and subsampled for phylogeny generation (see methods). The scalebar is provided in substitutions/site. Geographic maps are colored according to the sequence country metadata used for F13L analysis and do not represent the total geographic coverage of the MPXV clades.

| **F13L residue substitutions** | **IC_50_ (μM)** | **Fold change** |
| --- | --- | --- |
| K174N, N267D | 12 | 720 |
| H238Q | 0.54–0.6 | 28–34 |
| H238Q, N267D, A295E | 24 | 1,400 |
| H238Q, A288P, D294V, I372N | ≈5.2 | ≈290 |
| P243S, A288P, A290V | 0.56 | 32 |
| T245I, A290V | 0.17 | 10 |
| Y258C | 18 | 1,000 |
| N267D | 10–11 | 570–630 |
| N267D, A288P | 1.2–16 | 71–900 |
| N267D, A290V | 2.0 | 110 |
| N267D, D294V | 12 | 680 |
| N267D, A288P, A290V, D294V | >500 | >29,000 |
| N267D, A288P, A290V, A295E,L297ins | >500 | >29,000 |
| N267D, A288P, A290V, A295E,I372N | >500 | >29,000 |
| N267del | 1.5–4.0 | 85–230 |
| N267del, A290V | 0.13 | 7.5 |
| N267del, N267D, A295E | 2.9–18 | 160–1,000 |
| N267del, A288P, A295E | >500 | >29,000 |
| N267del, T289A, A295E | 0.26 | 15 |
| N267del, A290V, I372N | 3.1 | 180 |
| N267del, N267D, D294V, A295E | 2.5 | 140 |
| G277C | 2.3-40 | 95-800 |
| D283G | 7.1–7.3 | 404–420 |
| Y285H, I372N | 0.045 | 2.6 |
| A288P | 0.5 to >500 | 29 to >29,000 |
| A288P, I372N | >150 | >8,600 |
| A288P, A290V, D294V | 0.66 to >500 | 38 to >29,000 |
| A288P, A290V, L297ins | >500 | >29,000 |
| A288P, A290V, I372N | 15 | 880 |
| A288P, D294V, A295E | 1.4 | 83 |
| A288P, D294V, D301del | >500 | >29,000 |
| T289A | 0.078–0.14 | 3.7–7.8 |
| T289A, R291K | 1.7 | 98 |
| A290V | 0.17–43 | 10–2,500 |
| A290V, I372N | 30–32 | 1,700–1,800 |
| D294V | 0.23–1.4 | 13–78 |
| D294V, A295E | 1 | 59 |
| A295E | 2.0–3.3 | 110–190 |
| I372N | 0.04–>150 | 2.3 to >8600 |

#### **Table S1. Impact of amino acid mutations in F13L on MPXV resistance to Tecovirimat.**

The positions of amino acid mutations, either alone or in combination, conferring escape to Tecovirimat inhibition are indicated. The table was adapted from ^4^. The positions of amino acid mutations, either alone or in combination, conferring escape to Tecovirimat are indicated. Data are from references ^4^ ^5^ ^6^ ^7,8^.

|  |  |  | % within all MPXV sequences from GISAID | | | |  | % within its clade |
| --- | --- | --- | --- | --- | --- | --- | --- | --- |
| **Site** | **Mutation** | **Clade** | **Majority AA** | **Major AA Frequency** | **Minority AA** | **Minority AA Frequency** | **Genomes count** | **Minority AA Frequency** |
| **8** | P8S | 2b | P | 99.991% | S | 0.009% | 10968 | 0.009% |
| **29** | H29Y | 2b | H | 99.991% | Y | 0.009% | 10968 | 0.009% |
| **96** | S96R | 2b | S | 99.991% | R | 0.009% | 10968 | 0.009% |
| **100** | D100N | 2b | D | 99.927% | N | 0.073% | 10968 | 0.073% |
| **108** | I108M | 2b | I | 99.991% | M | 0.009% | 10968 | 0.009% |
| **109** | D109A | 2b | D | 99.982% | A | 0.009% | 10968 | 0.009% |
| **109** | D109H | 2b | D | 99.982% | H | 0.009% | 10968 | 0.009% |
| **110** | K110I | 2b | K | 99.991% | I | 0.009% | 10968 | 0.009% |
| **114** | V114A | 2b | V | 99.991% | A | 0.009% | 10968 | 0.009% |
| **172** | S172I | 2b | S | 99.991% | I | 0.009% | 10968 | 0.009% |
| **173** | A173S | 2b | A | 99.991% | S | 0.009% | 10968 | 0.009% |
| **174** | K174Q | 2b | K | 99.991% | Q | 0.009% | 10968 | 0.009% |
| **177** | W177L | 2b | W | 99.991% | L | 0.009% | 10968 | 0.009% |
| **184** | A184T | 2b | A | 99.770% | T | 0.230% | 10968 | 0.228% |
| **199** | I199V | 2b | I | 99.991% | V | 0.009% | 10968 | 0.009% |
| **215** | S215F | 2b | S | 99.991% | F | 0.009% | 10968 | 0.009% |
| **217** | D217N | 2b | D | 99.724% | N | 0.276% | 10968 | 0.274% |
| **229** | S229L | 2b | S | 99.991% | L | 0.009% | 10968 | 0.009% |
| **231** | K231N | 2b | K | 99.991% | N | 0.009% | 10968 | 0.009% |
| **238** | H238Q | 2b | H | 99.963% | Q | 0.037% | 10968 | 0.036% |
| **241** | I241T | 2b | I | 99.991% | T | 0.009% | 10968 | 0.009% |
| **243** | P243A | 2b | P | 99.917% | A | 0.083% | 10968 | 0.082% |
| **250** | N250S | 1a | N | 99.982% | S | 0.018% | 620 | 0.323% |
| **256** | D256N | 2b | D | 99.991% | N | 0.009% | 10968 | 0.009% |
| **267** | N267D | 2b | N | 99.760% | D | 0.230% | 10968 | 0.228% |
| **267** | N267S | 2b | N |  | S | 0.009% | 10968 | 0.009% |
| **278** | N278I | 2b | N | 99.982% | I | 0.009% | 10968 | 0.009% |
| **278** | N278S | 2b | N | 99.982% | S | 0.009% | 10968 | 0.009% |
| **280** | D280Y | 2b | D | 99.991% | Y | 0.009% | 10968 | 0.009% |
| **283** | D283G | 2b | D | 99.954% | G | 0.037% | 10968 | 0.036% |
| **283** | D283N | 2b | D |  | N | 0.009% | 10968 | 0.009% |
| **288** | A288P | 2b | A | 99.798% | P | 0.202% | 10968 | 0.201% |
| **290** | A290V | 2b | A | 99.743% | V | 0.257% | 10968 | 0.255% |
| **294** | D294N | 2b | D | 99.890% | N | 0.009% | 10968 | 0.009% |
| **294** | D294V | 2b | D |  | V | 0.101% | 10968 | 0.100% |
| **295** | A295E | 2b | A | 99.972% | E | 0.028% | 10968 | 0.027% |
| **296** | L296F | 2b | L | 99.991% | F | 0.009% | 10968 | 0.009% |
| **305** | K305R | 2b | K | 99.991% | R | 0.009% | 10968 | 0.009% |
| **307** | F307S | 2b | F | 99.991% | S | 0.009% | 10968 | 0.009% |
| **308** | T308A | 2b | T | 99.991% | A | 0.009% | 10968 | 0.009% |
| **335** | Y335C | 2b | Y | 99.991% | C | 0.009% | 10968 | 0.009% |
| **338** | H338L | 2b | H | 99.991% | L | 0.009% | 10968 | 0.009% |
| **352** | S352N | 1a | S | 99.991% | N | 0.009% | 620 | 0.161% |
| **353** | K353E | 2a | K | 91.351% | E | 8.649% | 48 | 4.167% |
| **353** | K353E | 2b | K |  | E |  | 10968 | 2.216% |
| **353** | K353E | 1a | K |  | E |  | 620 | 71.290% |
| **353** | K353E | 1b | K |  | E |  | 310 | 81.935% |
| **357** | I357M | 2b | I | 99.991% | M | 0.009% | 10968 | 0.009% |

#### **Table S2. Frequency of amino acid mutations in F13L associated with Tecovirimat resistance in all MPXV clades.**

The frequency of Tecovirimat resistance mutations within lineages based on the data available on the *GISAID EpiPox* database is indicated. The most frequent mutation in F13L, K353E is not located within the Tecovirimat binding site ^3^ and is not known to be associated with Tecovirimat resistance.

### Materials and Methods

#### **Cells and viruses**

Vero E6 and U2OS cells^9^ were grown in Dulbecco’s Modified Eagle Medium (DMEM, Gibco) supplemented with 10% fetal bovine serum (FBS, Gibco), 100 U/mL penicillin, and 100 μg/mL streptomycin (Gibco). Cells were maintained at 37°C and 5% CO_2_ for culture. Cells were routinely tested negative for mycoplasma. The MPXV isolates were amplified and titrated (plaque-forming units or PFU/mL) on Vero E6 cells.

The MPXV clade1a (LK isolate) strain was obtained from an infected patient in the Congo Basin. This virus was initially isolated on brain of newborn mice after intracerebral inoculation, then amplified on cell culture. The 3^rd^ cell passage of the virus was used ^10^.

The MPXV clade1b strain was isolated from a man in his mid-30s with no history of orthopoxvirus vaccination and traveling between Sweden and Africa in August 2024 ^1^. The isolate MpxV/PHAS-506/Passage- 03/SWE/2024_09_11, Clade 1b has been provided by the Public Health Agency of Sweden to improve the quality of diagnostics relevant for infectious disease control, treatment and/or other studies of relevance for public health. The 5^th^ cell passage of the virus was used.

The MPXV clade 2a strain (isolate Tai Nai Na, ref-SKU:010V-03360) was isolated from a skin lesion of an infected monkey (Sooty Mangabey) in Cote d’Ivoire in July 2012. The isolate, adapted on cell culture, was obtained from EVAg project (https://www.european-virus-archive.com/). The 5^th^ cell passage of the virus was used.

The MPXV clade 2b strain (MPXV/2022/FR/CMIP) was isolated from a pustular lesion of a 36-year-old French man who consulted at the Medical Center of Institut Pasteur (CMIP), in June 2022. The clinical specimen was inoculated on Vero E6 cells, whose supernatant was harvested after 3 days and tested positive for the presence of MPXV by PCR ^2^ ^10^. The titration of viral stocks was performed on U2OS cells ^2,11^. The 3^rd^ cell passage of the virus was used.

#### **Reagents**

Tecovirimat (BenchChem, catalogue number: B611274) was diluted at 10 mM in DMSO and stored at +4°C.

#### **MPXV inhibition assay**

U2OS cells were plated at 8 × 10^3^ cells per well in a μClear 96-well plate (Greiner Bio-One). The following day, cells were incubated with serial dilutions of Tecovirimat, starting 10μM. After four hours, the indicated MPXV strains were added to the cells in a BSL-3 facility. The viral inoculum was determined to obtain a non-saturating infection and similar number of foci of infected cells for each viral clade. Forty-eight hours later, cells were fixed for 30 min at room temperature (RT) with 4% paraformaldehyde (PFA, Electron Microscopy Sciences), washed and immunostained for MPXV antigens with rabbit polyclonal anti-VACV antibodies (PA1-7258, Invitrogen), and an Alexa Fluor 488-coupled goat anti-rabbit antibody (Invitrogen). Nuclei were stained with Hoechst (1:10,000; Invitrogen). Images were acquired with an Opera Phenix high-content confocal microscope (PerkinElmer). For each condition, infection was quantified by calculating the total area of MPXV-positive cells (MPXV^+^ area) and the nuclei were counted using the Harmony software (PerkinElmer). The percentage of infection inhibition was calculated from the MPXV^+^ area using the following formula:

$$100*\left[ 1-\left[ \frac{\left( {MPXV}^{+} area with Tecovirimat \right)-(mean area of 'non-infected^{'}controls)}{\left( mean area of 'no drug^{'} infected controls \right)-(mean area of 'non-infected^{'}controls)} \right] \right]$$

Inhibition activity of Tecovirimat was expressed as the IC50 (Half-maximal inhibitory concentration). IC50 were calculated based on an inhibitory dose-response curves with a variable slope model, using the percentage of inhibition at the different Tecovirimat concentrations.

#### **Biosafety**

All experiments with infectious MPXV were conducted under struct BSL3 conditions. Manipulations involving inactivated and non-inactivated MPXV were performed according to the French regulations on dual use pathogens.

#### **Sequence analysis**

Sequences were retrieved from the GISAID (https://www.gisaid.org ^12^) database on December 2, 2024 (Suppl tables 1,2). The F13L (VP37) nucleotide sequence was extracted from the MPXV reference sequence (NC_063383) and used to identify genomic coordinates for the F13L gene in all other MPXV genomes. In total, 11952 GISAID F13L sequences were utilized to compute the mutation frequencies. F13L nucleotide sequences were aligned using MAFFT v7.525. Incomplete codons i.e “nna” were considered missing data and converted to “nnn” to maintain an in-frame translation considering amino acid mutations. Mutations were identified independently within the two datasets to prevent sequence duplication and frequencies calculated in a clade-wise and clade specific manner.

For phylogeny construction, GISAID MPXV sequences were mapped to NC_063383.1 and subsampled by clade, year and country to retain 1 sequence per group using Augur (v27.0.0) ^13^. Phylogenies were visualized in R (v4.4.1) using the ggtree module (v3.12.0). Geographic maps were generated by extraction of the country metadata from the GISAID identifier string.

#### **Data availability**

All data supporting the findings of this study are available within the article or from the corresponding author upon reasonable request without any restrictions.

#### **Statistical analysis**

Figures were generated using Prism 9 (GraphPad Software). Statistical analysis was conducted using GraphPad Prism 9. Data are mean ±SD of triplicates from triplicates from one representative experiment, out of at least three independent experiments.

### Acknowledgements

The authors thank Klara Sondén (Department of Microbiology, Stockholm, Public Health Agency of Sweden) for sharing the MPXV clade 1b isolate, members of the Virus and Immunity Unit and other teams for discussion and help, Nathalie Aulner and the staff at the UtechS Photonic BioImaging (UPBI) core facility (Institut Pasteur), a member of the France BioImaging network, for image acquisition and analysis. This work has used the computational and storage services (Maestro cluster) provided by the IT department at Institut Pasteur, Paris.

We gratefully acknowledge all data contributors, i.e., the Authors and their Originating laboratories responsible for obtaining the specimens, and their Submitting laboratories for generating the genetic sequence and metadata and sharing via the GISAID Initiative, on which this research is based.

The MPXV clade 2a strain was obtained with the support of the European Virus Archive goes Global (EVAg) project that has received funding from the European Union’s Horizon 2020 research and innovative program under grant agreement N°653316.

The P.G.-C. lab is funded by Institut Pasteur, the National French Research Agency (ANR; ANR-22-CE11-0003). The E.S.-L. laboratory is funded by Institut Pasteur, the INCEPTION program (Investissements d’Avenir grant ANR-16-CONV-0005), the Ixcore foundation for research, the French Government’s Labex IBEID (ANR-10-LABX-62-IBEID), the HERA Project DURABLE (grant no 101102733) and the NIH PICREID (grant no U01AI151758). The O.S. lab is funded by Institut Pasteur, Fondation pour la Recherche Médicale (FRM), ANRS, the Vaccine Research Institute (VRI) (ANR-10-LABX-77), Labex IBEID (ANR-10-LABX-62-IBEID), the HERA projects DURABLE (grant 101102733) and LEAPS. JP is supported by DURABLE.

### Author contributions

*Experimental strategy design, experiments: JP, FGB, FP, OS.*

*Vital materials: QG, VC, JV, JCM, LD.*

*F13L sequences analysis and structure: JC, ESL, RV, PGC.*

*Manuscript writing and editing: JP, ESL, PGC, JCM, LD, OS.*
